# Impacts of thermal mismatches on chytrid fungus *Batrachochytrium dendrobatidis* prevalence are moderated by life stage, body size, elevation and latitude

**DOI:** 10.1101/473934

**Authors:** Jeremy M. Cohen, Taegan A. McMahon, Chloe Ramsay, Elizabeth A. Roznik, Erin L. Sauer, Scott Bessler, David J. Civitello, Bryan K. Delius, Neal Halstead, Sarah A. Knutie, Karena H. Nguyen, Nicole Ortega, Brittany Sears, Matthew D. Venesky, Suzanne Young, Jason R. Rohr

## Abstract

Global climate change is increasing the frequency of unpredictable weather conditions; however, it remains unclear how species-level and geographic factors, including body size and latitude, moderate impacts of unusually warm or cool temperatures on disease. Because larger hosts and lower-latitude hosts generally have slower acclimation times, we hypothesized that their disease susceptibility increases under “thermal mismatches”, or differences between baseline climate and the temperature during surveying. Here, we examined how thermal mismatches interact with body size, life stage, habitat, latitude, elevation, phylogeny, and IUCN conservation status to predict infection prevalence of the chytrid fungus *Batrachochytrium dendrobatidis* (*Bd*) in a global analysis of 38,967 amphibian hosts. As hypothesized, we found that the susceptibility of larger hosts and hosts from lower latitudes was strongly influenced by thermal mismatches. Furthermore, hosts of conservation concern are more susceptible than others following thermal mismatches, suggesting that thermal mismatches might have contributed to recent amphibian declines.

**Data Accessibility Statement:** Should the manuscript be accepted, the data supporting the results will be archived in an appropriate public repository such as Dryad or Figshare and the data DOI will be included at the end of the article.

## Introduction

In recent decades, climate change has increased global mean temperatures and the frequency of unusual and unpredictable weather conditions (Schär et al. 2004, Cai et al. 2014). Concurrently, infectious diseases of humans and wildlife have emerged at an accelerating pace, leading researchers to suggest a causal relationship between climate change and disease emergence (Jones et al. 2008, Liang and Gong 2017). However, the impacts of climate change on infectious disease are controversial (Harvell et al. 2002, Rohr et al. 2011, Altizer et al. 2013, Lafferty and Mordecai 2016), and several major knowledge gaps have persisted, preventing researchers from developing theory necessary to better understand the links between climate change and disease. First, although many laboratory experiments and field studies have addressed relationships between increasing mean temperatures and disease risk (e.g., Holmes et al. 2014, Paull and Johnson 2014, Ryan et al. 2017), relatively few studies have explored how variable and unusual conditions associated with climate change impact disease (but see Patz et al. 2005, Raffel et al. 2013, McIntyre et al. 2017). Second, researchers need more information about the types of hosts that will be most vulnerable to an interaction between disease and climate change (Rohr et al. 2011, Altizer et al. 2013). Third, there is consensus that most parasites and disease vectors are likely to expand poleward and towards higher elevations with climate change (Rohr et al. 2011, Altizer et al. 2013, Lafferty and Mordecai 2016), but some research also suggests that parasites will suffer at tropical latitudes and low elevations (Lafferty and Mordecai 2016). Studies that examine how species-level and geographic factors interact with variable weather conditions to cause disease outbreaks might allow researchers to predict which hosts will be most vulnerable to disease as climate change intensifies.

Recently, several frameworks have been proposed to explain how unusual climatic conditions could increase infectious disease outbreaks. One such framework, the *thermal mismatch hypothesis,* suggests that conditions that deviate from those typically experienced by hosts and parasites (i.e., thermal mismatches) will often favor parasites over hosts (Fig. 1) (Cohen et al. 2017, Cohen et al. in review) because smaller organisms generally have broader thermal breadths and acclimate or adapt to new conditions faster than larger organisms (Rohr et al. 2018). The *thermal mismatch hypothesis* makes two explicit assumptions: that hosts and parasites are generally adapted to typical conditions in their environments, and that they are limited by extreme conditions. However, the hypothesis is generally robust to violations of these assumptions (Fig. S1). The hypothesis suggests that hosts adapted to cooler environments should be most susceptible to parasites under unusually warm conditions, while hosts adapted to warmer environments should be most susceptible under unusually cool conditions. Thus, the hypothesis allows researchers to predict how host populations might fare when exposed to disease under different climate regimes. Although the physiological mechanisms underlying the *thermal mismatch hypothesis* are undetermined, it is likely that thermal stress reduces host immune response and general performance (Raffel et al. 2006) for longer periods of time than it reduces parasite performance, because parasites should acclimate and adapt to temperature shifts faster than their hosts (Raffel et al. 2013, Rohr et al. 2018).

**Figure 1.**
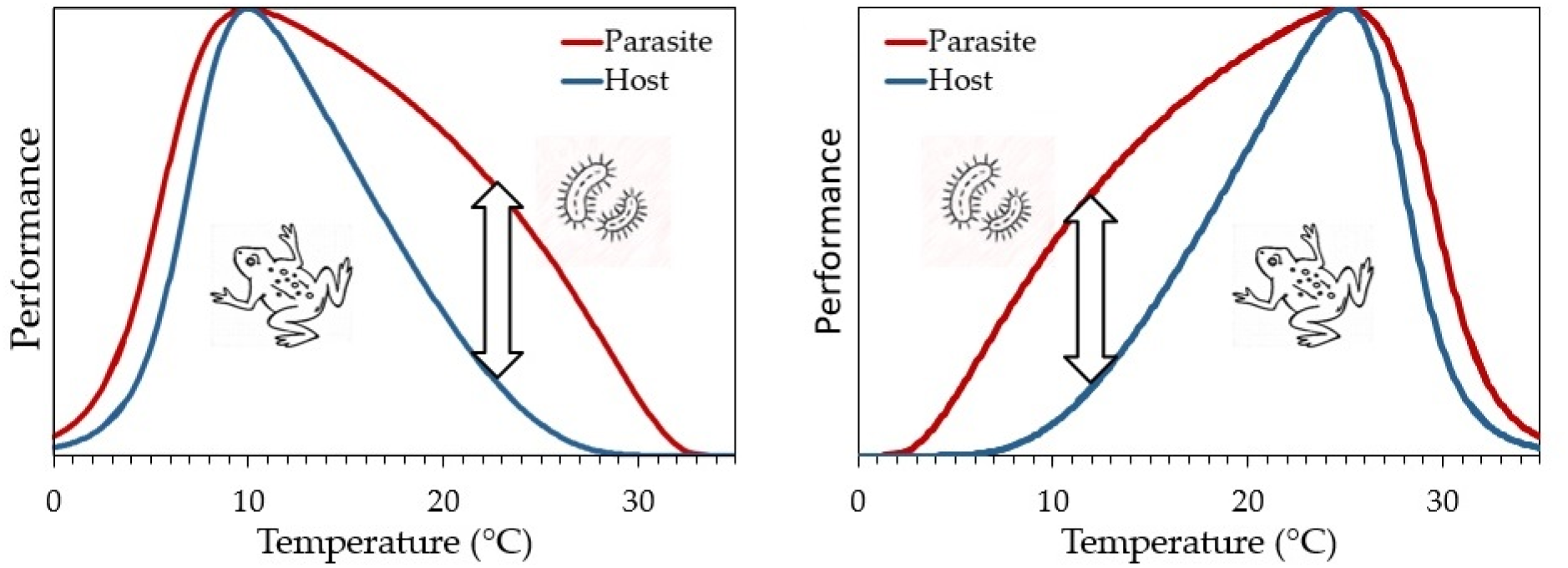
Conceptual figures representing the *thermal mismatch hypothesis* (reprinted from Cohen et al 2017). In isolation, small organisms, such as parasites (red lines), generally have broader thermal performance curves than larger organisms, such as hosts (blue lines). Parasite growth on hosts is likely to occur at temperatures where a parasite most outperforms its host (arrows), and not necessarily at the temperature at which a parasite performs best in isolation, providing a hypothesis for the thermal performance curve of a parasite growing on the host. For interacting cold-adapted hosts and parasites (a), parasite growth should be maximized at relatively warm temperatures, whereas for interacting warm-adapted hosts and parasites (b), parasite growth is predicted to be maximized at relatively cool temperatures. Previously published experimental and field studies support this hypothesis. For example, in experiments, Cohen et al. (2017) found that cool- and warm-preferring frogs had the highest infection loads of *Batrachochytrium dendropbatidis* (*Bd*) at warm and cool temperatures, respectively.

The *thermal mismatch hypothesis* has received support from several studies that used laboratory and field data to show that cold- and warm-adapted amphibians across the globe were most susceptible to the chytrid fungus, *Batrachochytrium dendrobatidis* (*Bd*), under warm and cool temperatures, respectively. Multiple temperature-dependent infection experiments in which amphibians did and did not have the ability to thermoregulate each demonstrated that cold-adapted amphibian host species experienced the greatest *Bd* growth at the warmest temperatures, while warm-adapted species experienced the greatest *Bd* growth at the coldest temperatures (Cohen et al. 2017, Sauer et al. 2018). A global analysis of amphibian populations that have been monitored for *Bd* similarly demonstrated that cold-adapted populations were most susceptible to disease under unusually warm conditions, and *vice versa* (Cohen et al. 2017; Fig. S2a). Furthermore, thermal mismatches outperformed other climatic factors predicting sharp declines and several extinctions experienced by the amphibian genus *Atelopus* in Latin America that are associated with *Bd* (Cohen et al. in review; Fig. S2b-c).

Despite the support for the *thermal mismatch hypothesis*, it is unclear which context-dependencies might influence the magnitude of the effects of thermal mismatches on host susceptibility to parasites and disease. For example, factors such as host geographic location (latitude and elevation), body size, habitat preference, life stage, and phylogeny can all influence host-parasite dynamics (Rohr et al. 2010, Johnson et al. 2012, Roznik and Alford 2015, Sears et al. 2015) and thus, might also moderate the influence of thermal mismatches on disease. Here, we evaluated how each of these factors affects the direction and magnitude of thermal mismatches—the differences between the historical mean temperature to which a host was exposed and the temperature during peak *Bd* prevalence—on historical, global records of *Bd* prevalence from 38,967 amphibians in 1,597 populations (Table S1). *Bd* is an ideal pathogen for testing hypotheses about thermal mismatches and climate change because it is easy to test for, well-monitored in a variety of locations worldwide, directly exposed to external air and water temperature on its ectothermic hosts, and has been associated with hundreds of worldwide amphibian declines (Skerratt et al. 2007). Additionally, there has been significant disagreement over whether *Bd* outbreaks occur during warm or cool periods and substantial geographic variation in the seasonal timing of *Bd* outbreaks (for example, Bosch et al. 2007, Kriger and Hero 2007). These issues can be addressed by the *thermal mismatch hypothesis* because it predicts that the effects of temperature on *Bd* should be very host-specific, depending on the temperatures to which a host is adapted.

Using our assembled database on global *Bd* prevalence, we tested the following hypotheses. Overall, we hypothesized a negative relationship between historical mean temperatures in the location of each population and the temperature during peak *Bd* prevalence because the *thermal mismatch hypothesis* predicts that disease should peak when warm-adapted species are subjected to cool conditions and *vice versa*. Second, we predicted that smaller host species might be less sensitive to an interaction between *Bd* and thermal mismatches than larger host species because smaller ectothermic host species should have lower thermal inertia and thus broader thermal breadths than larger hosts (Rohr et al. 2018, Rohr et al. in press), and the difference in their acclimation rates to new temperatures relative to that of *Bd* should be smaller than the same difference for larger hosts. Third, we predicted that tropical amphibian hosts should experience greater susceptibility to *Bd* following variable temperatures because the thermal breadths of these hosts should be narrower than those of temperate hosts as they typically experience less variation in temperature (Rohr and Raffel 2010). Meanwhile, the thermal breadths of *Bd* strains are typically much broader than those of hosts and do not vary much across latitude (Cohen et al. 2017, Voyles et al. 2017). Fourth, we predicted that hosts from high elevations would experience greater sensitivity to *Bd* under thermal mismatches than hosts from low elevations because they generally experience lower inter-annual and seasonal thermal variation (Brattstrom 1968, Navas 1996; also supported by our data - see results). Fifth, because water and soil temperatures fluctuate less than air temperatures (Lamoureux and Madison 1999, Lencioni 2004), animals inhabiting these habitats should be less vulnerable to *Bd* following thermal mismatches. Thus, we predicted that infection risk under thermal mismatches would be higher for (a) terrestrial species than aquatic species, (b) adults than larvae (which are more aquatic), and (c) Anurans than Caudates (which are more fossorial). Finally, we predicted that susceptibility to *Bd* following thermal mismatches would be associated with IUCN conservation status but did not have an *a priori* direction for this prediction.

## Materials and Methods

### Bd prevalence data

In July 2017, we searched Web of Science for the term *Batrachochytrium dendrobatidis*, producing 1,215 total results. Of these, we extracted data from 336 papers that reported *Bd* prevalence for at least one group of ≥5 amphibians of a given species at a specific location and time. The resulting dataset consisted of 38,967 animals from 1,597 sampling events, representing 440 species surveyed across 501 unique geographic locations around the globe (Fig. S3; Table S1). For each population, we extracted information on binomial species name, the number of infected and uninfected individuals, *Bd* prevalence, host developmental stage (larva, recent metamorph, and adult), month(s) of collection, elevation at host collection site, and geographic coordinates for host collection location directly from the paper. When not given in the paper, we identified coordinates on Google Maps based on papers’ descriptions, discarding records for which descriptions were not specific enough to identify sites to at least 1/10^th^ of one degree. We standardized all nomenclature according to the IUCN (International Union for Conservation of Nature; 2015). This dataset is an updated version of the *Bd* prevalence dataset used in Cohen et al (2017), with approximately 50% more data from an additional three years (2014-2017) of published results.

### Climate data and species-level traits

We extracted monthly mean temperature and precipitation data from the Hadley Climate Research Unit (Harris et al. 2016) for the location and specific month(s) that each amphibian population was swabbed in the field [*raster* package (Hijmans 2014), extract function; all data compilation and analyses were conducted in R 3.1.0 (2014)]. Monthly climate data were available at a resolution of 0.5°^2^ cells (approximately 50×50 km). Elevation for each species was collected from Bioclim rasters (Fick and Hijmans 2017) using the same methods. If sampling took place over the course of up to three consecutive months or less, and was not reported at the monthly level, then we averaged climate data across the sampling months. We did not use data collected over longer periods of time, because averaging climatic data across more than three months would have been inappropriate to seasonal analysis. We obtained long-term mean temperatures for each species’ natural geographic IUCN-defined range over 50 years from the amphibian species-level life history database compiled by Sodhi et al. (2008). We also updated the nomenclature to match the 2015 IUCN list (IUCN 2015). Hereafter, we refer to these 50-year mean temperature or precipitation values as the “historical mean temperature” or the “historical mean precipitation”. For each population identified to species, we attached species-level traits including average adult body size (snout-vent length, SVL, in mm), adult habitat (aquatic, arboreal, terrestrial, or mixed), IUCN threat classification, and taxonomic order and family, all sourced from the Sodhi et al. (2008) database, except for IUCN threat status (IUCN 2015).

### Statistical analyses

The *thermal mismatch hypothesis* predicts a negative relationship between historical mean temperature and temperature during peak *Bd* prevalence. Our first goal was to determine which temperatures were associated with peak *Bd* prevalence across different climates with different historical mean temperatures. First, we divided amphibian populations into overlapping bins, each comprising a range of 50-year mean environmental temperatures spanning 4°C (e.g., long-term mean of 10-14°C, 11-15°C, 12-16°C, etc. until 26-30°C). Within each of these bins, we then fit non-linear Weibull models with underlying binomial probability distributions to the relationship between mean temperature during the month of *Bd* testing and the number of infected and uninfected individuals at the population level. Weibull fits offer asymmetrical temperature performance curves that cannot contain negative y-axis values and produce a parameter estimate based on a negative log-likelihood function for the parameter of optimal temperature (also known as *T_opt_* and represented by *b* in the below equation; *a*, *c*, and *d* define the height, breadth, and skew of the curve, respectively) (Angilletta Jr 2006).

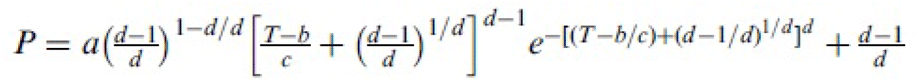

Thus, we generated estimates of *T_opt_* of *Bd* prevalence across a “moving window” of climate ranges. We did not fit a model to the output to confirm the linearity of the relationship between 50-year environmental temperature and temperature during sampling because the overlapping bins were not independent of one another.

We used generalized, binomial mixed-effects linear models (lme4 package, Bates et al. 2014; lmer function) to test for three-way interactions between a thermal mismatch effect (see below) and one of the following explanatory variables: life stage (adults or larvae), habitat type (terrestrial or aquatic; we excluded other types for a more direct comparison), order (Anura or Caudata only; we were unable to test order Apoda because of low sample size (*n*=6)), log-transformed mean elevation, absolute value of latitude, mean SVL, and IUCN classification (represented by two categories, least concern or a category including all species designated vulnerable, threatened, endangered, or extinct in the wild). Thermal mismatch effects were defined as negative statistical interactions between long-term mean temperature and temperature during the month(s) when *Bd* prevalence was tested at a given location. As we did not detect a thermal mismatch effect among larva, we restricted the data to adult amphibians when testing for effects of habitat type, order, elevation, latitude, body size, and IUCN status.

To isolate the effects of a given independent variable, we controlled for potential confounding variables as fixed effects. First, we controlled for latitude in models testing for effects of taxonomy, elevation, body size, habitat type, and IUCN status because latitudinal effects on *Bd* prevalence have been demonstrated repeatedly (Kriger et al. 2007, Becker and Zamudio 2011). Second, while testing for the three-way interaction among temperature during sampling, historical mean temperature, and latitude, we controlled for body size because body size and latitude covary and effects of latitude on acclimation are often dependent on body size (Rohr et al. 2018). Third, we attempted to control for habitat type when testing for a three-way interaction among temperature during sampling, historical mean temperature, and life stage to better understand whether aquatic habitats dictated the effects of thermal mismatches on larval amphibians, but the model would not converge, likely because there were too few terrestrial larvae in our dataset.

All models included study as a random effect to control for unknown confounding effects unique to a given sampling effort and/or potentially non-independent hosts. We wished to include spatial and phylogenetic correlation structures as random effects in our model, but this is not possible in a binomial mixed-model because unlike gaussian distributions, binomial distributions are defined by only one parameter which represents the mean, and thus do not have a parameter that exclusively defined the variance that can be modified by correlation structures. Still, we explored new methods to control for spatial structure of our data as a nested random effect, and although we have reservations about the reliability of these methods due to their novelty and the lack of available information about them, we report results of these models in the Supplement. To visualize our data, we created partial residual plots to isolate the effects of certain factors while controlling for all other factors in the statistical model that are not being displayed in the figure (generated with *visreg* package, *visreg* function). To further examine the influence of spatial effects in our primary *glmer* models, we plotted the residuals of some of our models in space (Fig. S4). Finally, to support our hypothesis that hosts from higher elevations are more susceptible to *Bd* following thermal mismatches than those from low elevations because they are adapted to less temperature variability, we conducted a Flinger-Killeen test for homogeneity of variances to test how the variance of mean temperature during sampling changed with elevation in our dataset.

## Results

For adult amphibians, our “moving window” approach revealed that there was a negative relationship between *T_opt_* of *Bd* prevalence and the historical mean temperature of host populations (Fig. 2). For species from climates with historical mean temperatures between 10 and 17°C, *Bd* prevalence peaked when temperatures were between 20-21°C, and for species from climates with historical temperatures between 21-28°C, *Bd* peaked between 13.5-15.5°C (Fig. 2).

**Figure 2.**
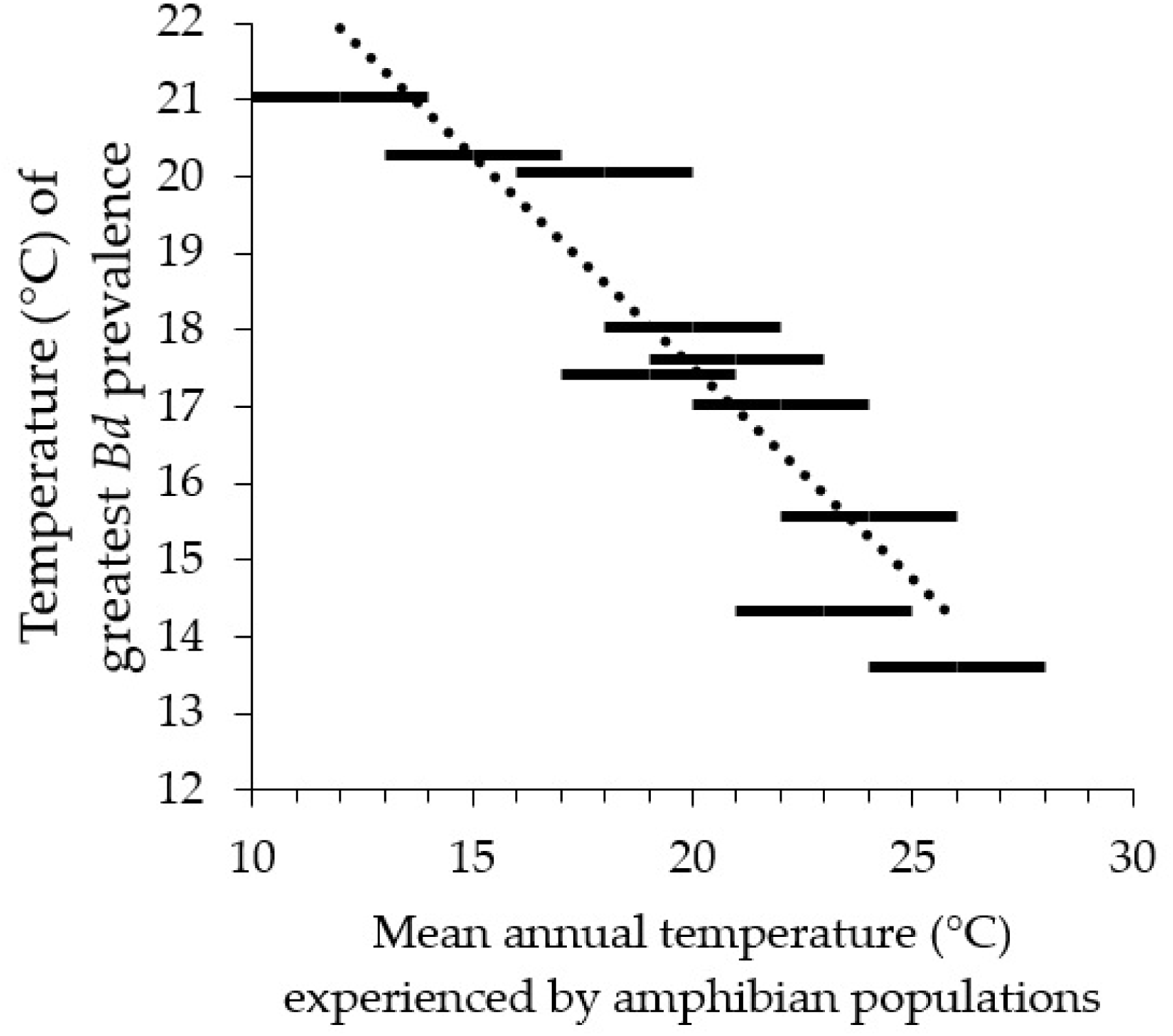
*Bd* prevalence peaks under cool conditions in typically warm climates and vice versa. To determine the temperatures during peak *Bd* prevalence across different climates with different historical mean temperatures, we fit non-linear Weibull models with underlying binomial probability distributions to the relationship between mean temperature during the month of *Bd* testing and *Bd* prevalence at the population level. We repeatedly fit such Weibull functions for overlapping subsets of amphibian populations, each comprising 4°C windows (e.g., 10-14°C, 11-15°C, 12-16°C, etc. until 26-30°C) of 50-year mean environmental temperatures, generating estimates of *T_opt_* of *Bd* prevalence across a “moving window” of climate types. Here, we extracted the Weibull model parameters describing the temperature at which *Bd* prevalence peaked and plotted these against 50-year mean annual temperatures experienced by amphibians in each window (bars). Some windows are missing as Weibull models would not fit these data due to low sample sizes.

In mixed-effects models, thermal mismatches, represented as negative interactions between the historical mean temperature and temperature during peak *Bd* prevalence, significantly predicted *Bd* prevalence in adults but were not related to *Bd* prevalence in larval amphibians (historical mean temperature *x* temperature during peak prevalence *x* life stage: β=0.0106, *p*<0.0001; Table S2; Figs. 3a-b). Further, amphibians considered “threatened” had higher *Bd* prevalence associated with thermal mismatches than species that were designated as “least concern” (historical mean temperature *x* temperature during peak prevalence *x* IUCN status: β=-0.0096, *p*<0.0001; Table S3; Figs. S3c-d). Amphibian species with larger adult body sizes had higher *Bd* prevalence associated with thermal mismatches than those with smaller adult body sizes (historical mean temperature *x* temperature during peak prevalence *x* body size; β=-0.000038, *p*=0.018; Table S4; Figs. 3e-f). The strength of thermal mismatch effects on *Bd* prevalence did not significantly depend on the habitat (terrestrial versus aquatic) of adult amphibians (historical mean temperature *x* temperature during peak prevalence *x* habitat; β=0.0101, *p*=0.163; Table S5; Fig. S5).

**Figure 3.**
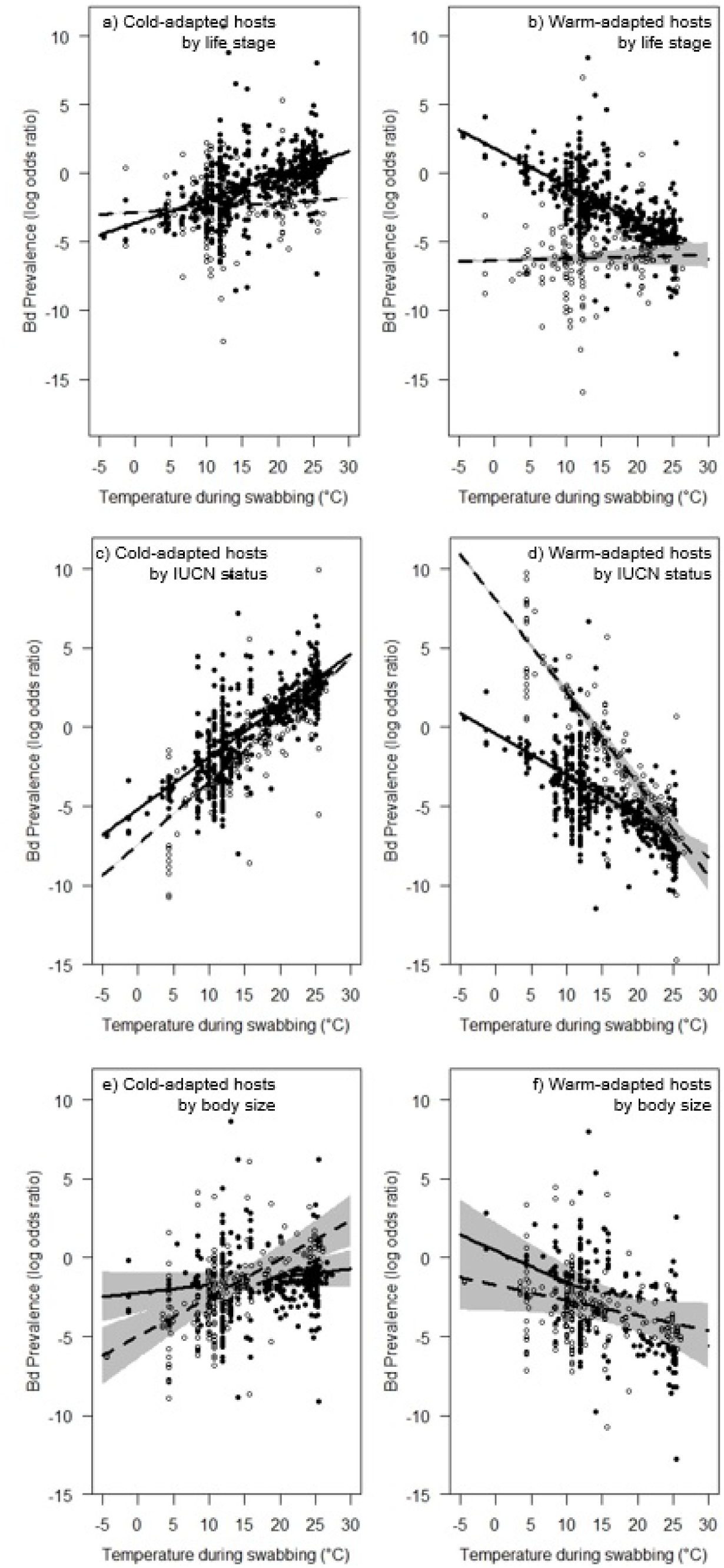
The effects of thermal mismatches on disease prevalence depend on host life stage, IUCN threat status, and adult body size. The partial residuals are from the binomial mixed-effects models shown in Tables S2, S3, and S4 (respectively) and display the significant three-way interactions among historical mean temperature, temperature during testing for *Batrachochytrium dendrobatidis* (*Bd*), and either life stage (a-b), IUCN threat status (c-d), or body size (e-f). Points represent individual amphibian populations tested for *Bd,* gray shading shows associated 95% confidence bands, solid lines and filled points represent adults, “least concern” populations and animals with smaller body sizes, respectively, while dashed lines and open points represent larvae, “threatened/” populations and larger body sizes, respectively. Left and right panels represent cold-adapted (a, c, e; 20^th^ percentile long-term mean temperatures centered on 9.90°C) and warm-adapted (b, d, f; 80^th^ percentile long-term mean temperatures centered on 24.01°C) hosts, respectively. The models suggest that adult – but not larval - amphibians from typically cooler climates experience higher *Bd* prevalence under warm conditions (a), while those from warmer climates experience high prevalence when conditions are cool (b). Among adults, species classified by IUCN as either vulnerable/threatened/endangered (combined into a single category) experience a stronger effect of thermal mismatches than those classified as “least concern” (c-d). Finally, amphibians from colder climates with larger body sizes (80^th^ percentile, ~113mm SVL) experienced greater *Bd* prevalence under warm conditions than smaller amphibians (e; 20^th^ percentile, ~33mm SVL), but showed no difference in temperature-dependent susceptibility in warm climates (f).

We found that the influence of thermal mismatches on *Bd* prevalence in adults depended significantly on geography and taxonomy. Thermal mismatches were associated with increased *Bd* prevalence more for amphibian populations at higher than lower elevations (controlling for latitude: historical mean temperature *x* temperature during peak prevalence *x* elevation; β=- 0.0051, *p*<0.0001; Table S6; Figs. 4a-b) and at lower than higher latitudes (when controlling for body size: historical mean temperature *x* temperature during peak prevalence *x* latitude; β=0.00042, *p*<0.0001; Table S7; Figs. 4c-d). A Flinger-Killeen test for homogeneity of variances confirmed that host species in our dataset did indeed experience greater temperature variability at lower elevations, providing a possible explanation why higher-elevation hosts may be less adapted to thermal mismatches (χ2=382.61, *p*<0.0001). Finally, thermal mismatches also increased *Bd* prevalence for Anura when controlling for latitude (historical mean temperature *x* temperature during peak prevalence *x* order; β=0.1427, *p*<0.0001; Table S8; Fig. S6), but thermal mismatches decreased *Bd* prevalence for Caudata.

**Figure 4.**
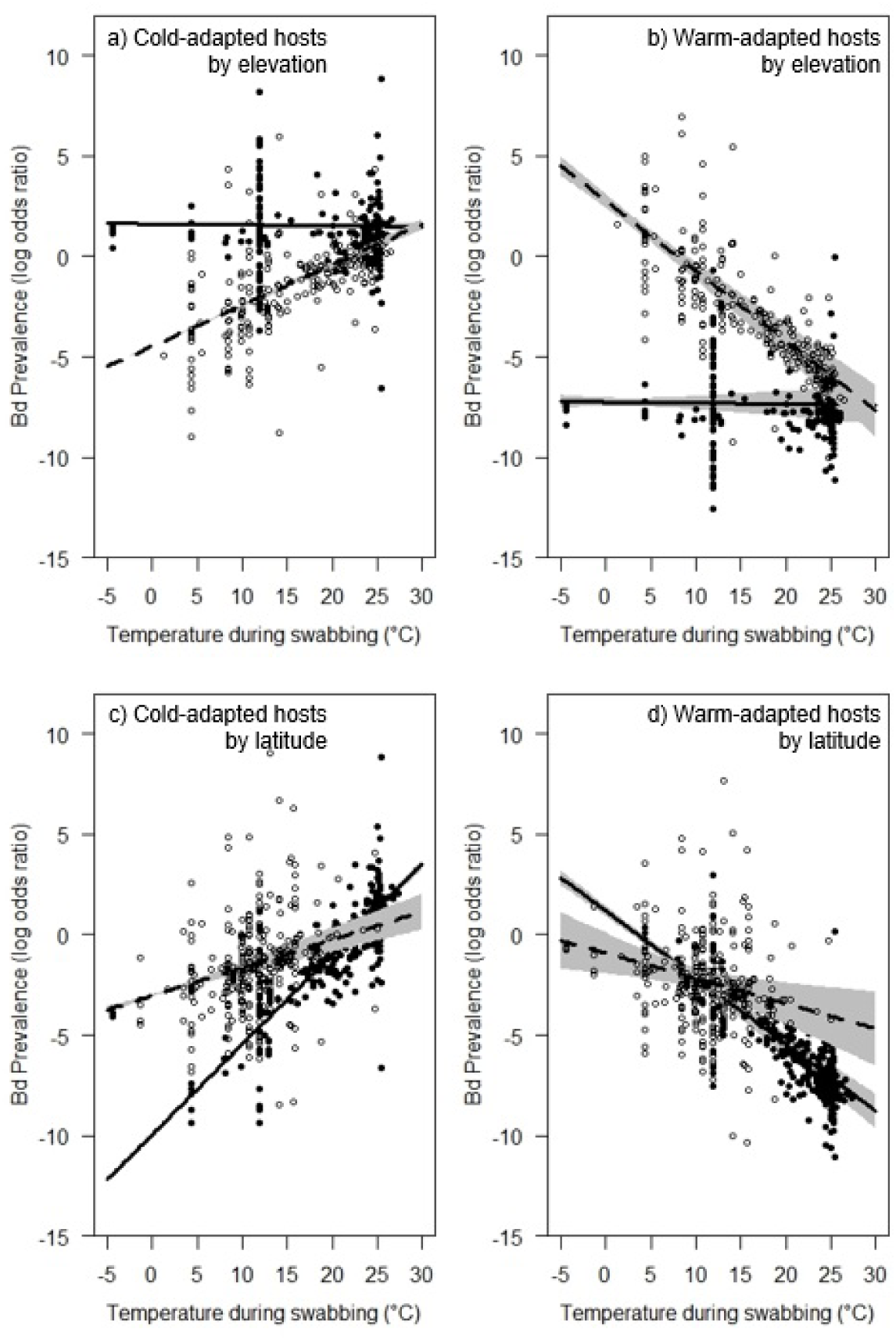
Interactions between thermal mismatches and disease prevalence depend on host body size. Partial residuals of *Batrachochytrium dendrobatidis* (*Bd*) prevalence are from the binomial mixed-effects models shown in Tables S6 and S7. Residuals display the significant three-way interaction among historical mean temperature, temperature during testing for *Bd* in a host population, and elevation (a-b) or latitude (c-d). Left and right panels represent cold-adapted (a, c; 20^th^ percentile long-term mean temperatures centered on 9.90°C) and warm-adapted (b, d; 80^th^ percentile long-term mean temperatures centered on 24.01°C) hosts respectively, points represent individual amphibian populations tested for *Bd* and gray shading shows associated 95% confidence bands. Solid lines and filled points represent low elevations (centered at ~118m) and latitudes (±12.0°), while dashed lines and open points represent high elevations (~1510m) and latitudes (±41.4°). The models suggest that at high elevations and low latitudes, amphibians from typically cooler climates experience higher *Bd* prevalence under warm conditions (a, c), while those from warmer climates experience high prevalence when conditions are cool (b, d). Meanwhile, populations from high latitudes show a reduced impact of thermal mismatches on their *Bd* prevalence (c-d), while populations from low elevations show no impact of thermal mismatches on *Bd* (a-b).

In models with additional spatial controls, interactions between thermal mismatches and host life stage, Order, and habitat remained unchanged (Tables S9-11). However, the interaction between thermal mismatches and elevation was no longer significant following the inclusion of spatial controls (Table S12). See Fig. S4 for plots of the model residuals in geographic space.

## Discussion

Our results support our hypothesis that hosts have increased susceptibility to infectious disease when experiencing unusual environmental temperatures and elucidate which hosts are likely to experience increased disease risk with climate change. As we predicted, adult amphibians, which are more often exposed to air temperature and thus experience greater temperature variability than aquatic larvae, demonstrated high susceptibility to *Bd* following thermal mismatches in their environments. Meanwhile, larval amphibians exhibited no such effect of thermal mismatches on infection prevalence, likely because water temperatures are more stable than air temperatures. However, adult amphibians experienced increased infection prevalence following thermal mismatches regardless of whether they were primarily aquatic or terrestrial, possibly because adults have more keratin on their skin and thus are more susceptible to *Bd* in general. An influence of thermal mismatches on infection was only apparent in Anurans, likely because Caudates are often fossorial and soil temperature fluctuates less than air temperature. Thus, groups of hosts that are typically directly exposed to air temperatures were more likely to experience an effect of thermal mismatches on their disease prevalence than hosts buffered from much of the variability in air temperature by either water or soil. Critically, we found that the susceptibility of host species considered “threatened” by the IUCN increases more following thermal mismatches than species of “least concern”, suggesting that thermal mismatches have played a role in recent amphibian declines linked to *Bd.* Alternatively, species’ traits that increase their vulnerability to the interaction between thermal mismatches and Bd might also independently increase their overall risk of extinction (e.g., lower genetic variability).

In our analyses, hosts with larger body sizes had a significantly higher *Bd* risk than smaller hosts following thermal mismatches, probably because they have relatively narrow thermal breadths compared to smaller-bodied hosts (Rohr et al. 2018, Rohr et al. in press). Additionally, larger hosts acclimate to changing temperatures at a slower pace than smaller hosts and thus are more susceptible to parasites for longer lengths of time after temperature shifts (Raffel et al. 2013). Our findings add to the growing body of evidence that larger organisms are experiencing greater costs of climate change across taxa and geographic locations (Estes et al. 2011, Lefort et al. 2015, Urban et al. 2017, Rohr et al. 2018), mainly because large organisms have comparatively greater thermal sensitivities and slower recovery times than small organisms. Systems with more drastic differences between host and parasite body sizes might be more severely impacted by the effects of thermal mismatch and disease because the differences between their thermal breadths are exaggerated relative to systems with smaller hosts (Fig. 1).

As predicted, thermal mismatches impacted susceptibility to *Bd* among hosts from lower latitudes to a greater extent than among hosts from higher latitudes, presumably because low-latitude hosts have narrower thermal breadths than high-latitude hosts (Rohr et al. 2018, Rohr et al. in press). In addition, we observed the reverse pattern for elevation; hosts experienced higher *Bd* prevalence after thermal mismatches at high but not low elevations despite our reasoning that high-elevation hosts would have narrower thermal breadths than low-elevation hosts and thus a reduced capability to deal with disease at unusual temperatures. Thus, hosts adapted to environments with typically lower thermal variation (low latitudes and high elevations) were less capable of coping with the combination of *Bd* and thermal mismatches.

Although this study addresses how thermal mismatches influence *Bd* across hosts and geographic locations, the effects of thermal mismatches across different host-parasite systems remain unclear (Rohr et al. 2011, Altizer et al. 2013). In general, we expect the influence of the *thermal mismatch hypothesis* to be more easily detectable among generalist parasites, because temperature-dependent infection patterns can be compared across cold- and warm-adapted hosts. We expect that ectothermic hosts are most likely to be affected by a combination of thermal mismatches and disease because they cannot internally regulate body temperatures.

Endoparasites of endotherms might be more advantaged by a combination of thermal mismatches and disease than ectoparasites of endotherms because these parasites are sheltered from direct exposure to fluctuating air temperatures. In addition, while we did not test for a latitude-disease relationship in polar hosts because there are no polar amphibians, polar organisms might suffer high disease risk following thermal mismatches because many have severely restricted thermal tolerances (Peck et al. 2004). Finally, although thermal mismatches were not detected among aquatic larval hosts in our study, other taxa with aquatic hosts that are highly sensitive to small changes in temperature (e.g., corals; Rowan 2004) could be highly vulnerable to disease under thermal mismatches.

Our results suggest that the effects of thermal mismatches on host-parasite interactions are more complex than previously thought and are highly dependent on host species, life stage, and geographic location. In accordance with previous evidence suggesting that climate change should have greater adverse effects on larger than smaller organisms, we found that larger hosts might have increased susceptibility to infectious disease after experiencing unusual temperatures, likely because of their often-narrower thermal breadths than both smaller hosts and parasites.

Furthermore, we found that hosts experience higher *Bd* prevalence following thermal mismatches at lower than higher latitudes and at higher than lower elevations, likely because hosts adapted to these areas are less accustomed to temperature variation. Although bacterial, viral, and helminth diseases have traditionally been predicted to become deadlier with climate change (Harvell et al. 2002, Lafferty 2009), our results provide a framework to explain how climate change could sometimes exacerbate fungal disease outbreaks under highly variable conditions for hosts and contrast with previous suggestions that fungal diseases might be reduced by climate change (Fisher et al. 2012). Our results provide evidence that hosts adapted to cooler environments will likely suffer increased parasitism as climate change increases the frequency of relatively warm conditions, whereas hosts from warmer environments will experience less parasitism because climate change is decreasing the number and length of cool periods.

## Acknowledgments

Thanks to A. Diesmos and D. Bickford for allowing us to use their dataset describing life history traits of amphibian species. Thanks to O. Santiago and S. Spencer for providing thoughts on the manuscript and to A. Carey for assistance assembling Table S1. Funds were provided by grants to J.R.R. from the National Science Foundation (EF-1241889 and DEB-1518681), the National Institutes of Health (R01GM109499, R01TW010286-01), the US Department of Agriculture (2009-35102-0543), and the US Environmental Protection Agency (CAREER 83518801), and to T.A.M. from the Faculty Development Dana and Delo grants at the University of Tampa.

Author contributions
J.M.C. and J.R.R. conceived ideas, J.M.C. and T.A.M. oversaw construction of database, all authors extracted data from the literature to build the dataset, J.M.C. and E.A.R. standardized nomenclature and added species traits to the dataset, J.M.C. acqired climate and species-level data, J.M.C. conducted statistical analyses, J.M.C. and J.R.R. wrote the paper, and all authors contributed to manuscript revisions.

